# Nucleosome turnover is sufficient to establish varied histone methylation states

**DOI:** 10.1101/256321

**Authors:** Emma J. Chory, Joseph P. Calarco, Nathaniel A. Hathaway, Oliver Bell, Dana S. Neel, Gerald R. Crabtree

## Abstract

Transcription-dependent methylation of histone H3 at lysine 79 (H3K79) is evolutionarily conserved from yeast to mammals, critical for normal development and frequently deregulated by genetic recombination in Mixed Lineage Leukemia. Although this histone modification is associated with gene activity, little is known about the cellular mechanisms of H3K79 methylation regulation. Because no H3K79 demethylase has been discovered, the mechanism of its removal remains unclear. Utilizing chemical-induced-proximity to control histone methylation *in vivo* we show that the dynamics of methylation state (mono, di, tri-methylation) is genome-context specific. Further, Monte Carlo simulations coupling systems of kinetic reactions with histone turnover rates, suggest that nucleosome turnover is sufficient to establish varied genome-wide methylation states without active demethylation.

---

The histone methylation state of the epigenome is mediated through antagonizing enzymatic processes of “writing” and “erasing” histone modifications. This is often mediated by “reader” protein domains which can recognize specific histone marks to manifest unique biologic outcomes (*1*). Methylation of Lysine 79 on Histone 3 (H3K79) is mediated by the multi-subunit disruptor of telomeric silencing (DOT)-complex. Despite being evolutionarily conserved and implicated in the development of Mixed Lineage Leukemia (MLL) (*2*), regulation by the DOT-complex remains poorly understood at a mechanistic level. Mono-, di-, and tri-H3K79 methylation, (H3K79me1, H3K79me2, H3K79me3) marks are solely mediated by the DOT1L methyltransferase (*3*) which is localized to unmodified H3K27 sites through a direct interaction between the octapeptide motif-leucine zipper (OMLZ) domain of AF10/AF17 (*4*). The complex is recruited specifically to active sites of the genome through a direct interaction of AF9 with H3K9Ac (*5*). Subunits of the DOT-complex are commonly translocated in recurrent MLL-rearranged leukemia and result in dysregulation of the homeobox (HOX) gene cluster which is critical for persistence of the disease (*6*, *7*). H3K79me1, H3K79me2, and H3K79me3 likely play divergent biologic roles in both normal and leukemic contexts, as the these marks are associated with active transcription to varying degrees (*8*, *9*) and efficient tri-methylation of H3K79 requires monoubiquitination of H2BK123 (*10*). Despite the development of many potent chemical probes targeting the DOT1L methyltransferase (*11*, *12*), it remains unclear if and how the H3K79 methylation mark is removed, and how varied methylation states are established, as no H3K79 demethylase “eraser” has been identified to date.

In this study, we tethered the OMLZ domain of the DOT-complex subunit AF10 to a synthetic DNA binding domain (ZF-DBD) and used chemical-induced proximity (CIP) (*13*, *14*) to selectively target H3K79 methylation at endogenous, un-methylated genes *in vivo*. In doing so, we observed the kinetics of mono-, di-, and tri-methylation in real-time within diverse genomic contexts and chromatin substrates and determined that nucleosome turnover is sufficient to establish local methylation states, in the absence of active demethylation.

We generated two distinct, novel murine embryonic stem cell lines (mESCs) containing an array of DNA binding domains (12xZFHD1) upstream of either the *Ascl1* or the *Hbb-y* transcription start site (TSS) along with an in-frame nuclear enhanced green fluorescent protein (EGFP) replacing the first exon (Fig. 1A) (*15*-*17*). Two fusion proteins, Zinc-finger/FKBP and AF10(OMLZ domain)-FRB were stably expressed in the mESC recruitment lines by lentiviral transduction to permit rapid deposition of H3K79 methylmarks using small molecule (Rapamycin)-mediated recruitment (Fig. 1A). *Ascl1* and *Hbb-y* were specifically targeted as they represent two distinct types of H3K79me-deficient genomic sites in mESCs with differing chromatin landscapes. Insertion of the DNA binding arrays at *Hbb-y* represents a “naïve” target locus and enables monitoring chromatin dynamics independently of other endogenous regulatory factors (e.g., transcription factors, remodeling factors, etc.). In contrast, insertion at a bivalent gene (*Ascl1*) allows monitoring H3K79 dynamics at a natural MLL target, but in the absence of active transcription and pre-existing H3K79 methylation (Fig. 1B) (compared to at a highly-transcribed pluripotency factor such as Oct4). These two genetic loci served as distinct chromatin substrates which allowed us to interrogate H3K79me dynamics *in vivo* and in real-time. Rapamycin-induced deposition of the H3K79me3 mark was observed at both recruitment sites, yet to varying degrees (Fig. 1B). At the *Hbb-y* locus, a H3K79me3 domain was slowly and gradually formed over a period of seven days (Fig. 1C), while at *Ascl1* the maximum enrichment was reached by 2 days (Fig. 1D). We confirmed that the placement of the mark is catalytic, by inhibition with the Dot1L inhibitor, EPZ-04777 (Fig. S1). However, recruitment of the DOT-complex and subsequent placement of the H3K79me3 mark did not result in expression of the EGFP reporter at the *Ascl1* locus (Fig. S2). While it may seem surprising that H3K79 methylation is insufficient to activate a heavily H3K4me3-decorated bivalent gene in this context, this result does not rule out the possibility that H3K79me is sufficient to initiate transcriptional activation at other loci. Expression of the *Ascl1* transcription factor is heavily reliant on a positive feedback mechanism, and as such, was purposefully knocked-out in the mESC reporter line to prevent spontaneous neural lineage differentiation upon induction (*18*).

**Figure 1.**
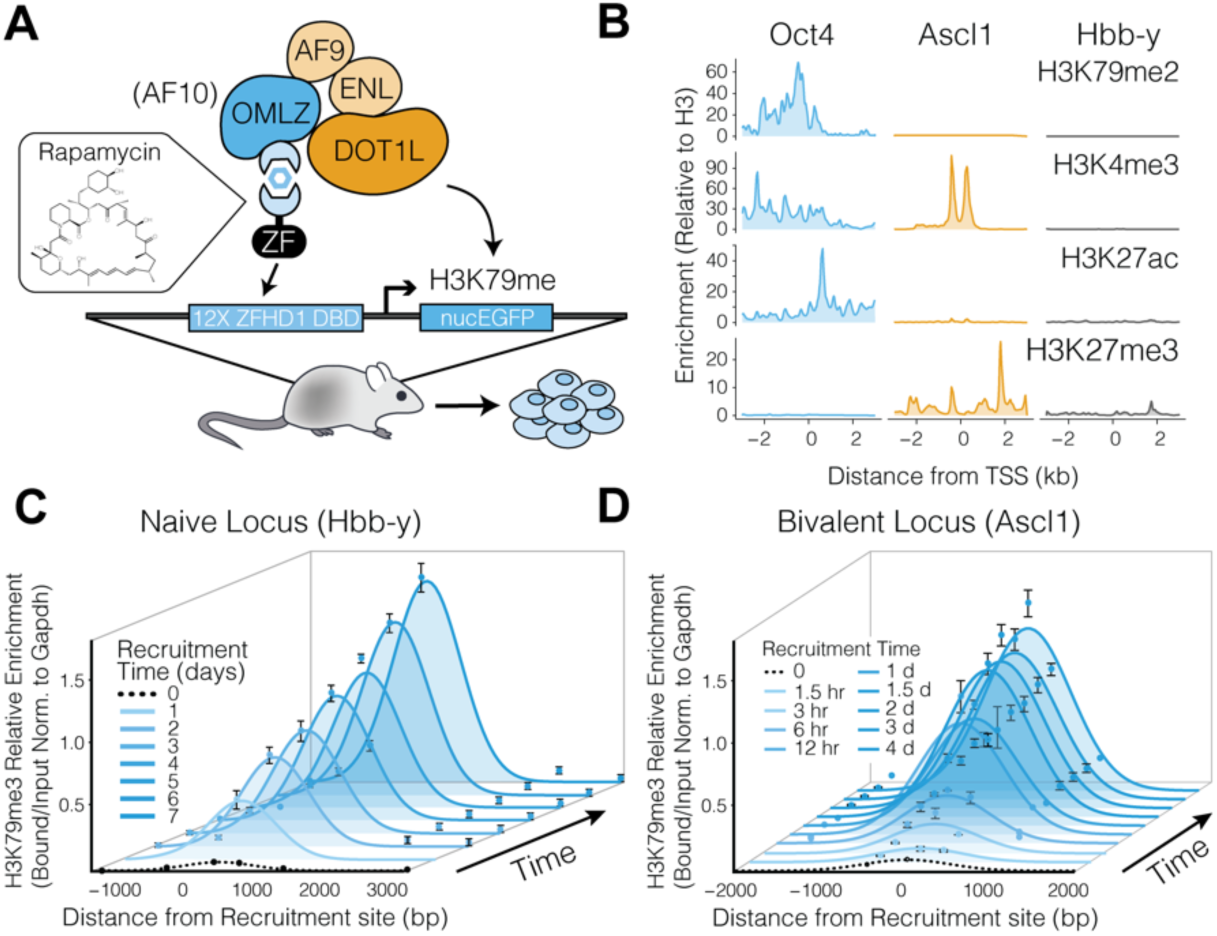
Design of and placement of H3K79me3 in mESC targeting lines. **A)** The *CIP* knock-in lines contain a single modified allele harboring two arrays of DNA binding sites (12XZFHD1 and 5XGal4) in the promoter region upstream of an in-frame EGFP reporter. **(B)** Distribution of histone modifications at *Hbb-y* and *Ascl1* in murine ES cells (*21*, *32*), compared to the active *Oct4* site reveals the distinct, un-methylated H3K79me2 chromatin substrates for induced modulation. **(C)** ChIP analysis reveals dynamic changes of H3K79me3 chromatin modifications at the *Hbb-y* locus, over the course of 7 days of Rapamycin-mediated recruitment of Dot1L. **(D)** ChIP analysis reveals dynamic changes of H3K79me3 chromatin modifications at the *Ascl1* locus, over the course of 4 days of Rapamycin-mediated recruitment of Dot1L, with a maximum occurring at < 2 days of recruitment. Error bars represent n=3 independent experiments.

To determine the source of the distinctive methylation dynamics at the *Ascl1* and *Hbb-y* loci, we examined the rates of mono-, di-and tri-methylation over the course of 7 days. We find that while H3K79me3 continually builds at *Hbb-y* over 7 days, mono-methylation is rapidly acquired and reaches a maximum by 12 hours but then steadily decreases, with H3K79me2 exhibiting a slightly delayed progression, reaching a maximum at 2 days and then similarly decreasing (Fig. 2A). On the contrary, at *Ascl1* the opposite is observed. H3K79me1, H3K79me2, and H3K79me3 are each rapidly induced, with mono-methyl establishing the highest enrichment by ~12 hours, followed by di-, then tri-methylation (Fig. 2B). Interestingly, despite *Ascl1 and Hbb-y* having similar pre-equilibrium methylation profiles (1 day of Rap-mediated DOT-complex recruitment) (Fig. 2C), at equilibrium (7-days of recruitment) *Ascl1* is most heavily decorated by H3K79me1 while *Hbb-y* is predominately H3K79 tri-methylated (Fig. 2C).

**Figure 2.**
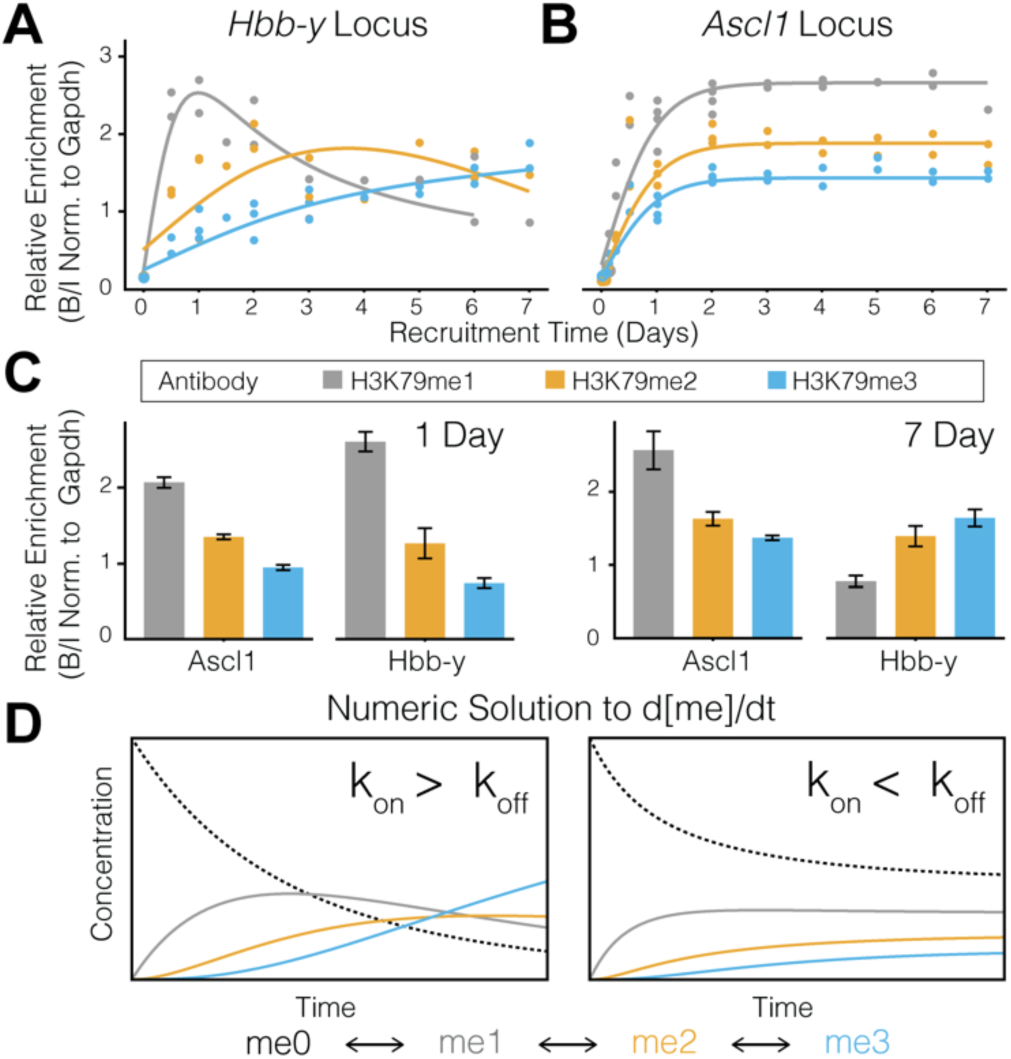
Kinetics of H379me1, me2, and me3 at various substrates. **A)** ChIP analysis at the recruitment site of the *Hbb-y* locus for H3K79me1, H3K79me2, and H3K79me3 over the course of 7 days (n=3 experiments). **(B)** ChIP analysis at the recruitment site of the *Ascl1* locus for H3K79me1, H3K79me2, and H3K79me3 over the course of 7 days (n=3 experiments). **(C)** Average H3K79me1, H3K79me2, and H3K79me3 at pre-equilibrium (1day), and post-equilibrium (7day) time points at Ascl1 and Hbb-y. **(D)** Analytical solutions to the system of ordinary differential equations describing the reactions from me0 ⇌ me1 ⇌ me2 ⇌ me3 for the conditions k_on_ ≫ k_off_ (left) and k_on_ ≪ k_off_ (right).

Intrigued by the opposing dynamics of these systems, we compared the respective ChIP kinetic profiles to the analytical solution of the system of ordinary differential equations (ODEs) describing the processive-kinetic reactions of the transition from:

> me0 ⇌ me1 ⇌ me2 ⇌ me3(Fig. S3).

The H3K79me dynamics profile observed at *Hbb-y* is consistent with the solution to a case where the forward rate of methylation (k_on_) is much greater than the reverse rate of methylation (k_off_), while the converse is observed for *Ascl1* (Fig. 2D). Thus, given that the protein level and rate of recruitment (i.e. concentration of small molecule) is constant, at *Ascl1* H3K79me is likely being removed by an active process.

Since Dot1L is the only H3K79 methyltransferase and to-date no histone demethylase has been identified, we sought to ascertain which regulatory factors may contribute to the differing dynamics observed at the two substrates. Similar to actively transcribed genes, bivalent sites undergo active nucleosome turnover and chromatin remodeling in order to maintain a poised state at promoters of developmentally important genes (*19*, *20*). This is thought to be mediated by the deposition of H3.3 through a direct interaction with PRC2, which results in the incorporation of newly synthesized histones (*21*). To estimate the relative rates of nucleosome turnover at *Hbb-y* and *Ascl1*, we examined previously published H3.3 ChIP-seq and CATCH-IT datasets (*21*-*23*) (Table S1) in mESCs at the respective genes and observed that the rate of nucleosome exchange at *Ascl1* is approximately ~100 fold greater than at *Hbb-y* in mESCs (Fig. 3A). Utilizing these respective nucleosome turnover rates, we developed a Monte Carlo model which simulates the kinetics of a methyltransferase catalyzing the reaction of an unmodified nucleosome transitioning from me0 → me1 → me2 → me3, based on the probability (e^-kt^ _on_) of the reaction occurring within a given timeframe. We added the condition that the modified nucleosome may also move to the right or left within an array of nucleosomes at either a high or low rate (e^-kt^_right,left_), and importantly held the methylation rate constant (at k_on_ ≫ k_off_) (Fig. 3B). As anticipated, at the site of recruitment, the results of the simulation of a “low turnover” rate, closely resemble the solution to the ODEs with the condition of k_on_ ≫ k_off_. Additionally, minimal methylation domain propagation upstream or downstream was observed (Fig. 3C). Conversely, the results of the simulation with high turnover rate (100x) results in local equilibrium concentrations in which me1 > me2 > me3. However, unlike the low turnover rate condition, the profile propagates upstream and downstream establishing a dominant mono-methyl domain that spreads bi-directionally over several kb (Fig. 3C).

**Figure 3.**
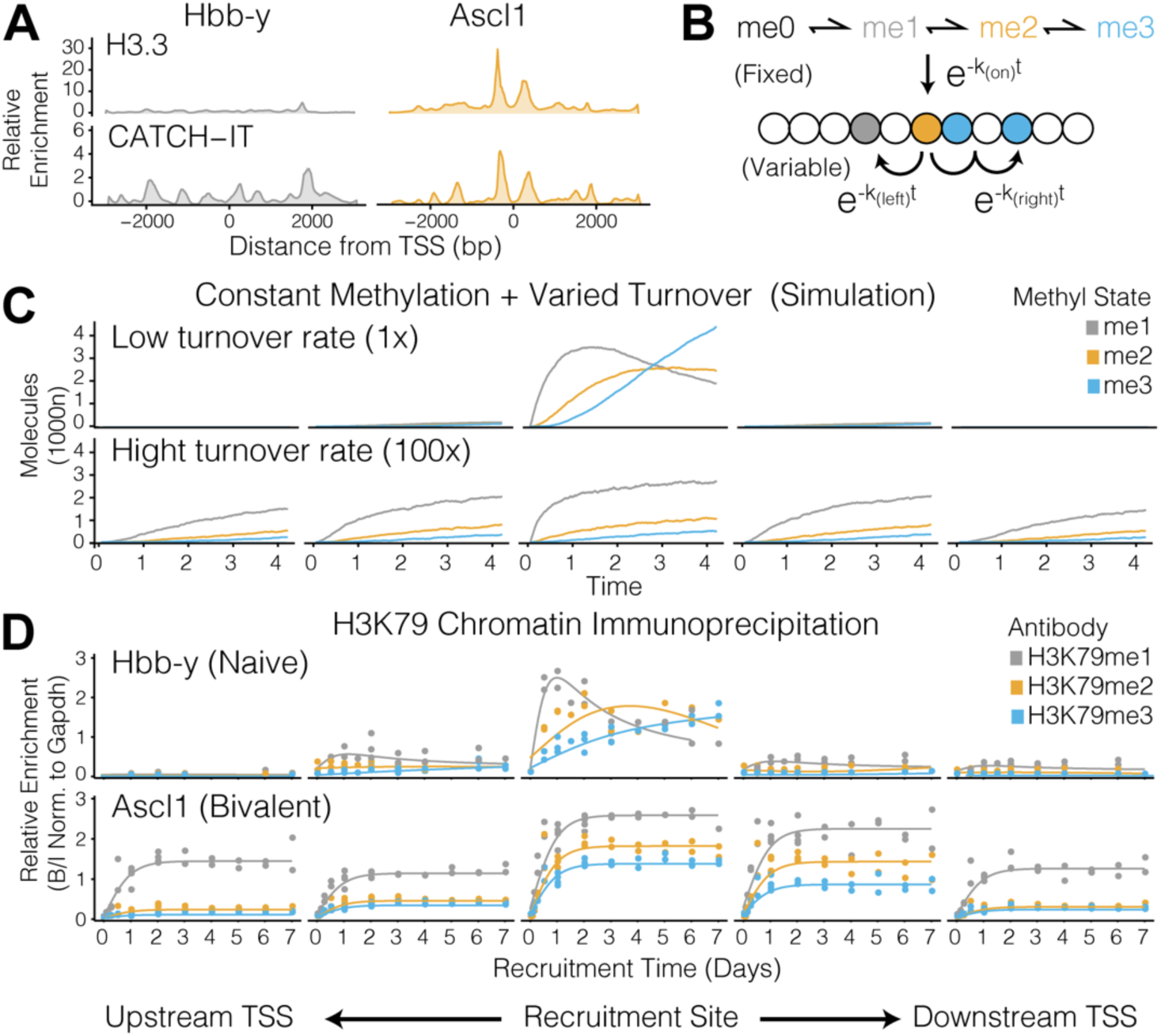
Model of methylation and nucleosome turnover supports varied substrate dynamics. **A)** Distribution of histone modifications (H3.3) and nucleosome incorporation rates (CATCH-IT) at *Hbb-y* and *Ascl1* in mES cells (*21*, *22*), indicative of nucleosome turnover rates **(B)** Monte Carlo simulation of methylation and nucleosome turnover considers the various chromatin recruitment substrates as a one-dimensional beads-on-a-string. Poisson processes of H3K79 methylation nucleation site and nucleosome turnover along the lattice are included. Me1, me2, and me3 are nucleated at the recruitment site at a rate of k_on_ ≫ k_off_ and turnover of the nucleosomes is equally likely across the lattice and occurs at rates of k_right_, k_left_. **(C)** Results of the Monte Carlo simulations showing −2 to +2 nucleosomes, for the conditions of “low nucleosome turnover” (top) and “high nucleosome turnover” (bottom) demonstrate local accumulation of H3K79me3 marks, and bidirectional propagation of H3K79me1 dominant domains, respectively. **(D)** ChIP analysis reveals dynamic changes of H3K79me1, H3K79me2 and H3K79me3 localized chromatin modifications at the *Hbb-y* locus (top) and propagation of modifications at the *Ascl1* locus (bottom). Each data point represents a single immunoprecipitation, at a given rapamycin recruitment time point, in each cell line, for n=3 independent experiments.

When we compare the Monte Carlo profiles of “high” and “low turnover” rates to the time-dependent levels of H3K79me1, H3K79me2, and H3K79me3 upstream and downstream of the TSS at both *Ascl1* and *Hbb-y*, it becomes immediately evident that the *in vivo* methylation dynamics we measured by ChIP recapitulate the predicted methylation profiles obtained from the simulation of high and low nucleosome turnover, respectively. The variable spreading of methylation and subsequent establishment of dominant H3K79me1 versus H3K79me3 domains most likely reflects that a central feature of establishing the distinct methylation states is differential nucleosome turnover, even in the absence of active demethylation. This model is supported by the observation that H3K79me1 levels are highly correlated with levels of active transcription (*6*), since actively transcribed genes undergo the most robust exchange of nucleosomes. Further, given that narrow AF9 peaks are primarily localized near the TSS of transcribed genes despite the establishment of broad H3K79me domains (Fig. S4) (*24*, *25*), a nucleosome turnover model is favored over oligomerization-based processive propagation (*13*, *26*). While our results certainly do not rule out the possibility of an undiscovered H3K79 “eraser” or undermine the critical functions of other histone demethylases, our model suggests that nucleosome turnover can be sufficient to establish unique methylation states across a variety of genomic contexts, without invoking enzymatic demethylation.

Extensive genome-wide ChIP-sequencing has been performed to characterize the phenotypic addiction of MLL-rearranged leukemias to aberrant H3K79me (*4*, *6*, *7*, *27*, *28*). As such, we sought to investigate whether a nucleosome turnover driven model may prove relevant in understanding the fundamental mechanisms driving this disease. To determine the relative methylation states genome-wide, at varying chromatin substrates, we explored previously published ChIP-seq and RNA-seq data sets in both mature mouse B cells (*29*), and MLL-AF9 leukemias derived from lineage–Sca-1+c-Kit+(LSK)-cells from mouse bone marrow (*6*) (Table S1). To determine whether nucleosome turnover plays a role in the establishment of the aberrant methylation which contributes to oncogenesis, we specifically looked at “high expressing”, “low expressing”, “bivalent”, and “polycomb repressed” genes in both healthy, mature B cells and the MLL-AF9 leukemias. Although advances have been made to improve the quantitative nature of ChIP-sequencing by adding spike-in controls (*30*), challenges remain to make direct, quantitative comparisons across samples with varying antibodies. To address this issue, we began by normalizing the log10 transformed average read density at each transcription start site (+/-10kb) within each dataset (Fig. S5), and used unbiased k-means clustering of the datasets (Table S1) to identify genes that are characteristic of “high”, “low”, “bivalent”, or “repressed” sites. For healthy B cells, “high expressing” genes were characterized by average RNA levels (GRO-seq), robust levels of H3K4me3 and H3K27Ac, moderate levels of H3K4me1, and no H3K27me3. “Low expressing” genes were identified similarly, however with less H3K27Ac. “Bivalent” genes were identified by moderate-to-high levels of both H3K27me3 and H3K4me3, while “Polycomb repressed” genes were characterized as having only H3K27me3 (Fig. 4A, Fig. S6). In MLL-AF9 transformed leukemias, average RNA levels of each gene and H3K27me3 levels were used to identify each cluster, along with H3K79-methylation levels (Fig. 4A, Fig. S7). Both stimulated and resting B cells along with AF10(-/-) and AF10(+/+) transformed leukemias were included to improve clustering efficiency of relevant genetic loci (Fig. S6-7). As predicted by the results of our CIP-mediated recruitment experiments and turnover simulations, we observe that in both mature B cells and MLL-AF9 transformed leukemias, “low expressing” and “high expressing” genomic loci are characterized by H3K79me1 > H3K79me2 > H3K79me3, which propagates over the gene body (Fig. 4B-C). This characterization is consistent with our nucleosome turnover model (Fig. S8), as transcribed genes exhibit high levels of nucleosome turnover in the direction of transcription. We further observe that while there is little H3K79me present at polycomb repressed sites, the dominant state is H3K79me3, which is consistent with the “low turnover” model and coincides with repressed genes (Fig. 4B-C).

**Figure 4.**
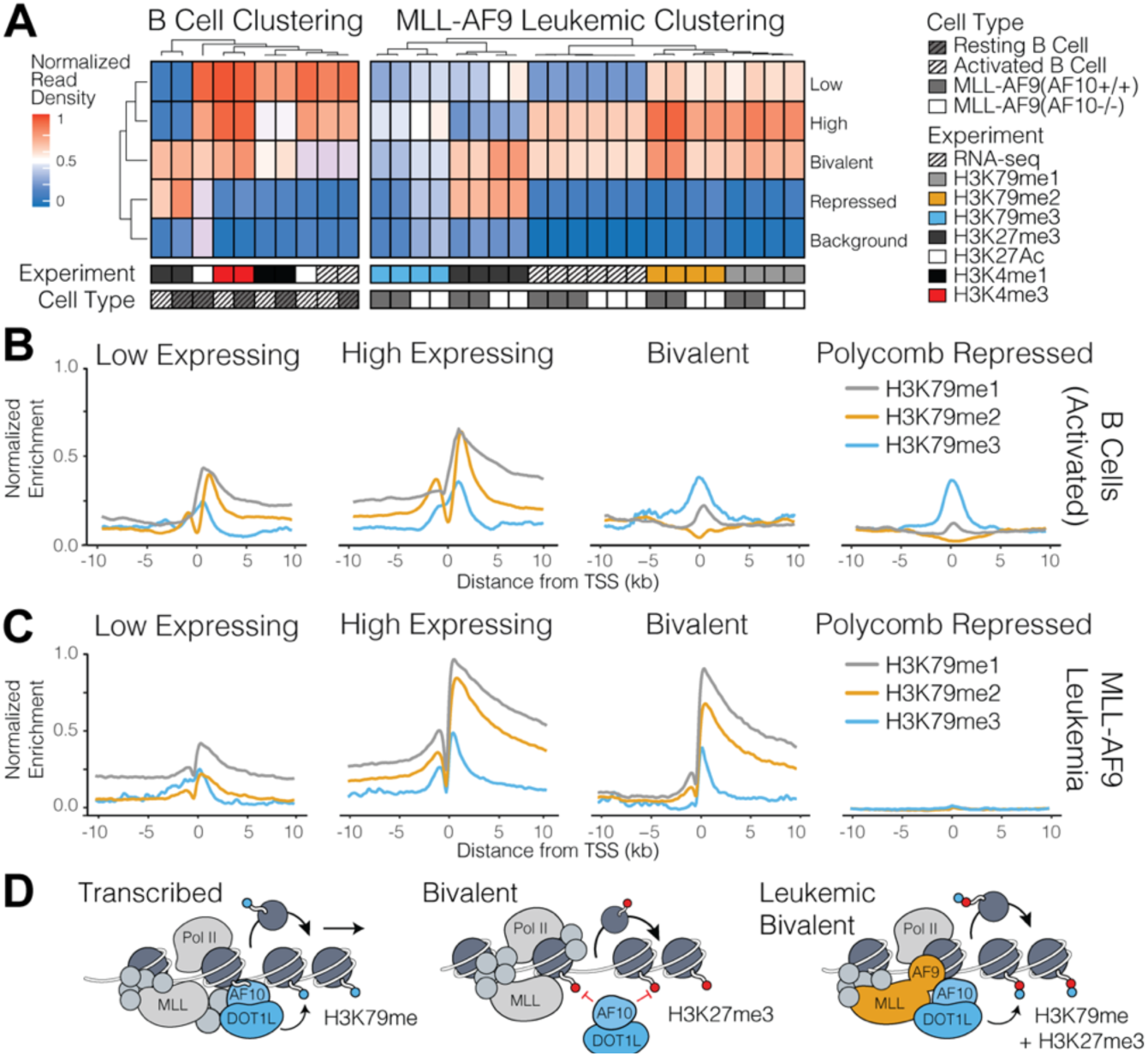
Model of methylation and nucleosome turnover supports varied substrate dynamics. **(A)** Clustering of genomic domains in mature B cells (left) (*29*) and MLL-AF9 leukemias (right) (*6*) by *k*-means, with *k* = 5. Clustering domains were separated into “high expressing”, “low expressing”, “bivalent”, “polycomb repressed” and “background” genes. **(B)** Meta-analysis of the averaged ChIP-seq signal for H3K79me1, H3K79me2, H3K79me3, and H3K27me3 at four sets of genes across a −10 kb to +10 kb genomic region around the transcriptional start sites of mature, activated B cells. **(C)** Meta-analysis of the averaged ChIP-seq signal for H3K79me1, H3K79me2, H3K79me3, and H3K27me3 at four sets of genes across a −10 kb to +10 kb genomic region around the transcriptional start sites of AF10+ MLL-AF9 transformed murine leukemia. **(D)** Model for nucleosome turnover establishment of the methylation state of H3K79.

Interestingly, specifically in mature B cells, “bivalent” sites resemble the H3K79 methylation profile of polycomb repressed sites, akin to the basal methylation state we observed at *Ascl1* in mESCs prior to CIP-mediated recruitment of AF10 (Fig. 4B). Conversely, in MLL-AF9 leukemias, the methylation profile at “bivalent” sites starkly resembles that of an actively transcribed gene, despite being moderately decorated with H3K27me3 (Fig. 4C). This methylation profile is consistent with the profile at *Ascl1* in mESCs following rapamycin-induced recruitment. Thus, in leukemia, the aberrant translocation of DOT-subunits with the MLL protein, and subsequent deposition of K79 methylation at bivalent genes likely establishes discrete H3K79me1 domains as a result of high levels of nucleosome turnover present at these regions. Given that in a normal context, the presence of H3K27me3 would obstruct the binding of AF10 to bivalent genes, (*4*), in MLL-AF9 leukemias, under our model, these highly turned-over sites would likely be hyper-sensitive to mis-targeting. We attribute this sensitivity to the high levels of nucleosome turnover and lack of active enzymatic demethylation which establish dominant, activating H3K79me1 peaks at these poised genomic regions.

Establishing histone methylation states requires an orchestra of readers, writers, and erasers to signal varied biologic outputs (*31*). Although the DOT-complex is associated with increased levels of transcription, and translocations are strongly implicated in the pathogenesis of Mixed Lineage Leukemias, little is understood about how the varied methylation states of H3K79 are established or removed. While histone demethylases are presently considered the main opposition force in many other methylation models, the analyses presented here suggest that nucleosome turnover is likely an active contributor to establishing mono-, di-, or tri-methylation domains. Moreover, the correlation between high levels of active transcription and H3K79me1 appear to be simply explained by processive kinetics, which predict that in the absence of demethylation and presence of nucleosome turnover, mono-methylation predominates. A nucleosome turnover model supports both the presence of small H3K79me3 peaks in the absence of turnover, and may account for the vast H3K79me1 domains that are established at poised H3K4me3 targets in MLL-rearranged leukemias. Whether H3K79me can be removed by a yet-to-be-discovered demethylase remains unclear, however our model provides evolutionary rationale supporting the lack of a demethylase for such a conserved and critical histone regulator. Further, nucleosome turnover and polycomb may regulate the H3K79 methylation state in opposition (Fig. 4D), with H3K27me3 simply excluding Dot1L from targeting genomic regions in which nucleosomes are being turned over but not actively transcribed. While the methylation dynamics presented here are simple, and alternative hypothesis such as varied binding affinities or demethylation remain possible, the H3K79 methylation state appears to be characterized by processive-first order kinetics, governed by first-principles, and largely influenced by the rate of nucleosome turnover.

## Acknowledgements

We wish to dedicate this manuscript to the lasting memory of Joseph P Calarco, who was a brilliant scientist, great friend and incisive colleague and mentor. We thank Dr. James Bradner and Jun Qi for providing EPZ-04777 along with their support and guidance, and Dirk Schübeler for providing the targeting Hbb-y cell line. We thank the Crabtree lab, J.G. Kirkland, S.M. Braun, B.Z. Stanton, CM Weber, W Wenderski, C. Hodges, NM Plugis, and A.J. Spakowitz for helpful discussions. E.J.C. was supported by an NSF Graduate Research Fellowship and the Ruth L. Kirschstein National Research Service Award (F31 CA203228-02). This study was supported by the Howard Hughes Medical Institute, NIH 5R01CA163915-04, the NIH Javits Neuroscience Investigator Award R37 NS046789-12, the CDMRP Breast Cancer Research Breakthrough Award, and funding from the Simons Foundation Autism Research Initiative.

